# Coordinated regulation of CB1 cannabinoid receptors and anandamide metabolism stabilizes network activity during homeostatic downscaling

**DOI:** 10.1101/2021.05.21.445170

**Authors:** Michael Ye, Sarah K. Monroe, Sean M. Gay, Michael L. Armstrong, Diane E. Youngstrom, Fabio L. Urbina, Stephanie L. Gupton, Nichole Reisdorph, Graham H. Diering

## Abstract

Neurons express overlapping homeostatic mechanisms to regulate synaptic function and network properties in response to perturbations of neuronal activity. Endocannabinoids (eCBs) are bioactive lipids synthesized in the post-synaptic compartments to regulate synaptic transmission, plasticity, and neuronal excitability primarily through retrograde activation of pre- synaptic cannabinoid receptor type 1 (CB1). The eCB system is well-situated to regulate neuronal network properties and coordinate pre- and post-synaptic activity. However, the role of the eCB system in homeostatic adaptations to neuronal hyperactivity is unknown. To address this issue, we used western blot and targeted lipidomics to measure adaptations in eCB system to bicuculline (BCC)-induced chronic hyperexcitation in mature (>DIV21) cultured rat cortical neurons, and used multielectrode array recording and live-cell imaging of glutamate dynamics to test the effects of pharmacological manipulations of eCB on network activities. We show that BCC-induced chronic hyperexcitation triggers homeostatic downscaling and a coordinated adaptation to enhance tonic eCB signaling. Hyperexcitation triggers first the downregulation of fatty acid amide hydrolase (FAAH), the lipase that degrades the eCB anandamide, then an accumulation of anandamide and related metabolites, and finally a delayed upregulation of surface and total CB1. Additionally, we show that BCC-induced downregulation of surface AMPA-type glutamate receptors (AMPARs) and upregulation of CB1 occur through independent mechanisms. Finally, we show that endocannabinoids support baseline network activities before and after downscaling and is engaged to suppress network activities during adaptation to hyperexcitation. We discuss the implications of our findings in the context of downscaling and homeostatic regulation of oscillatory network activities.

**Significance statement:** Neurons are remarkably resilient to perturbations in network activities thanks to the expression of overlapping homeostatic adaptations. In response to network hyperactivity or silencing, neurons respond through regulating excitatory and inhibitory post-synaptic neurotransmitter receptors density, probability of pre-synaptic neurotransmitter release, and/or membrane excitability. The endocannabinoid system is a prominent signaling pathway at many synapses that is known to be involved in multiple forms of short- and long-term synaptic plasticity. Here we find that components of the endocannabinoid system are upregulated in response to chronic hyperexcitation of cultured cortical neurons, and that endocannabinoid signaling is required to maintain network activity but also suppresses network events during hyperexcitation. This work supports a novel tonic homeostatic function for the endocannabinoid system in neurons.

## Introduction

Neurons express overlapping homeostatic mechanisms to maintain excitability and network properties in response to changes in synaptic inputs. Homeostatic scaling is a well described bidirectional phenomenon that broadly controls the strength of excitatory and inhibitory synapses through regulation of pre- and post-synaptic functions (Turrigiano, 2008; Turrigiano et al., 1998). In response to prolonged hyperexcitation or silencing, homeostatic mechanisms are engaged in cortical and hippocampal neurons to restore activity levels to a set point through compensatory regulation of post-synaptic AMPARs density (Diering et al., 2014; O’Brien et al., 1998), GABAergic transmission (Turrigiano, 2008), and/or pre-synaptic vesicle release (De Gois et al., 2005; Wierenga et al., 2006). The expression locus of homeostatic mechanisms varies depending on the neuronal types and maturation, with additional pre- synaptic mechanisms emerging ∼3 weeks *in vitro* in cultured primary cortical/hippocampal neurons (De Gois et al., 2005; Turrigiano, 2008; Wierenga et al., 2006). Whereas many molecular mechanisms have been described underlying pre- and post-synaptic homeostatic adaptations, it is not entirely clear how pre- and post-synaptic glutamatergic and GABAergic plasticity mechanisms are coordinated to maintain stable network activity at baseline and in response to network hyperexcitation.

Endocannabinoids (eCBs) are bioactive lipids that regulate neurotransmitter release and synaptic plasticity, largely through activation of pre-synaptic Cannabinoid receptor type 1 (CB1) (Castillo et al., 2012). CB1 is broadly expressed in many neuron types and is known to regulate glutamatergic and GABAergic transmission, primarily through a Gα_i_-coupled suppression of synaptic vesicle release (Castillo et al., 2012; Di Marzo, 2018). CB1 is also known to regulate cortical up-states, a prominent mode of slow oscillatory network activity seen in isolated cortical slices or dissociated cultures in vitro, or during non-rapid eye movement (NREM) sleep in vivo (Johnson and Buonomano, 2007; Pava et al., 2014; Saberi-Moghadam et al., 2018; Steriade et al., 1993a). Two major eCBs, 2-arachidonyl glycerol (2-AG), and arachidonoyl ethanolamide (AEA, also called anandamide) may act as agonists for CB1, but they are synthesized and degraded through completely non-overlapping sets of enzymes, suggesting that they may serve distinct functions and undergo separate regulation (Di Marzo, 2018). Unlike other neuromodulators, eCBs are not stored in vesicles for release, but are synthesized “on demand” from the catabolism of post-synaptic phospholipids in an activity-dependent manner. The eCB system constitutes an important retrograde signaling mechanism whereby activity of post- synaptic neurons can regulate pre-synaptic transmitter release (Castillo et al., 2012). Therefore, the eCB system is ideally situated to coordinate the activities of pre- and post-synaptic plasticity and regulate network activity during homeostatic scaling.

In the current study we have observed that in relatively mature cortical cultures, CB1 protein expression and cell-surface targeting are upregulated in response to chronic (48hr) hyper-excitation induced by the GABA_A_ antagonist bicuculline (BCC). This adaptation is coordinated with a downregulation of fatty acid amide hydrolase (FAAH), and consequent accumulation of AEA and related N-acylethanolamides (NAEs) to enhance tonic CB1-AEA signaling. We show that the well-described downregulation of surface AMPARs during downscaling is independent of CB1 signaling. Using multi-electrode array recordings and live-cell imaging of a fluorescent glutamate neurotransmitter reporter (iGluSnFr) (Marvin et al., 2018), we observe highly synchronized up-state-like oscillatory network activities in dissociated neuron culture (Pava et al., 2014; Saberi-Moghadam et al., 2018; Steriade et al., 1993a).

Consistent with a previous report (Pava et al., 2014), we show that CB1 signaling is required to maintain this up-state-like network activity, both under control conditions and following BCC- induced downscaling. Additionally, we show that enhancing CB1 further suppresses network activities during hyperexcitation. We propose that coordinated regulation of eCB metabolism and signaling is needed to adapt and maintain network activity during the response to network hyperexcitation.

## Materials and Methods

### Primary Neuron Culture

Primary dissociated cortical neurons were prepared from E18 Sprague-Dawley rats (https://www.criver.com/products-services/find-model/cd-sd-igs-rat?region=3611) as previously described [redacted for double-blind review]. Briefly, dissociated cortical neurons were plated onto tissue culture dishes coated with 1mg/mL poly-L-lysine (PLL, Sigma-Aldrich). Neurons were maintained in at 37℃/5% CO_2_, and fed twice a week in glial conditioned neurobasal media (Gibco) supplemented with 2% B27, 1% horse serum, 2mM GlutaMAX and 100U/mL penicillin/streptomycin (Gibco). For biochemistry experiments, neurons were plated at 600,000 cells/well into 6-well tissue culture plates (Falcon). For lipidomics experiments, neurons were plated at 5,000,000 cells/plate into 10cm tissue culture dishes (Corning). For live-cell imaging, neurons were plated at 120,000 cells/well into 4-well chambered cover-glass (Cellvis). For MEA recording, neurons were plated at 120,000 cells/wells into 24-well Cytoview MEA plates (Axion Biosystems).

### Drug Treatment

Drugs were prepared as 1000x stocks and stored at -20℃. To induce homeostatic scaling, neurons were treated with 20μM bicuculline methobromide (BCC) (Tocris, 20mM stock in water) or 1μM tetrodotoxin citrate (TTX) (Abcam, 1mM stock in water) for 48h. In some live-cell imaging experiments, 20μM BCC or 1μM TTX was applied acutely for 10-60min. For Figure 2, neurons were treated with 20μM BCC for 4, 12, 24 or 48h to monitor the process of homeostatic down-scaling. For Figure 3, neurons were treated for 48h with a combination of 1μM MTEP hydrochloride (Tocris, 1mM stock in DMSO) and 100nM JNJ 16259685 (Tocris, 100μM stock in DMSO) to block mGluR1/5 signaling. For Figure 4, neurons were treated for 48h with 100nM PF3845 (Cayman, 100μM stock in DMSO) or 2.5μM PF3845 (Cayman, 2.5mM stock in DMSO) to inhibit FAAH and to elevate AEA and related NAEs. For Figure 5, neurons were treated for 6h with 50nM AM251 (Cayman, 50μM stock in DMSO), or for 10-60min with 500nM AM251 (Cayman, 500μM stock in DMSO) to block CB1 signaling. For Extended Fig 5-1, neurons were treated for 10-60min with 50μM D-AP5 (Cayman, 50mM stock in water) to block NMDA signaling.

**Figure 1.**
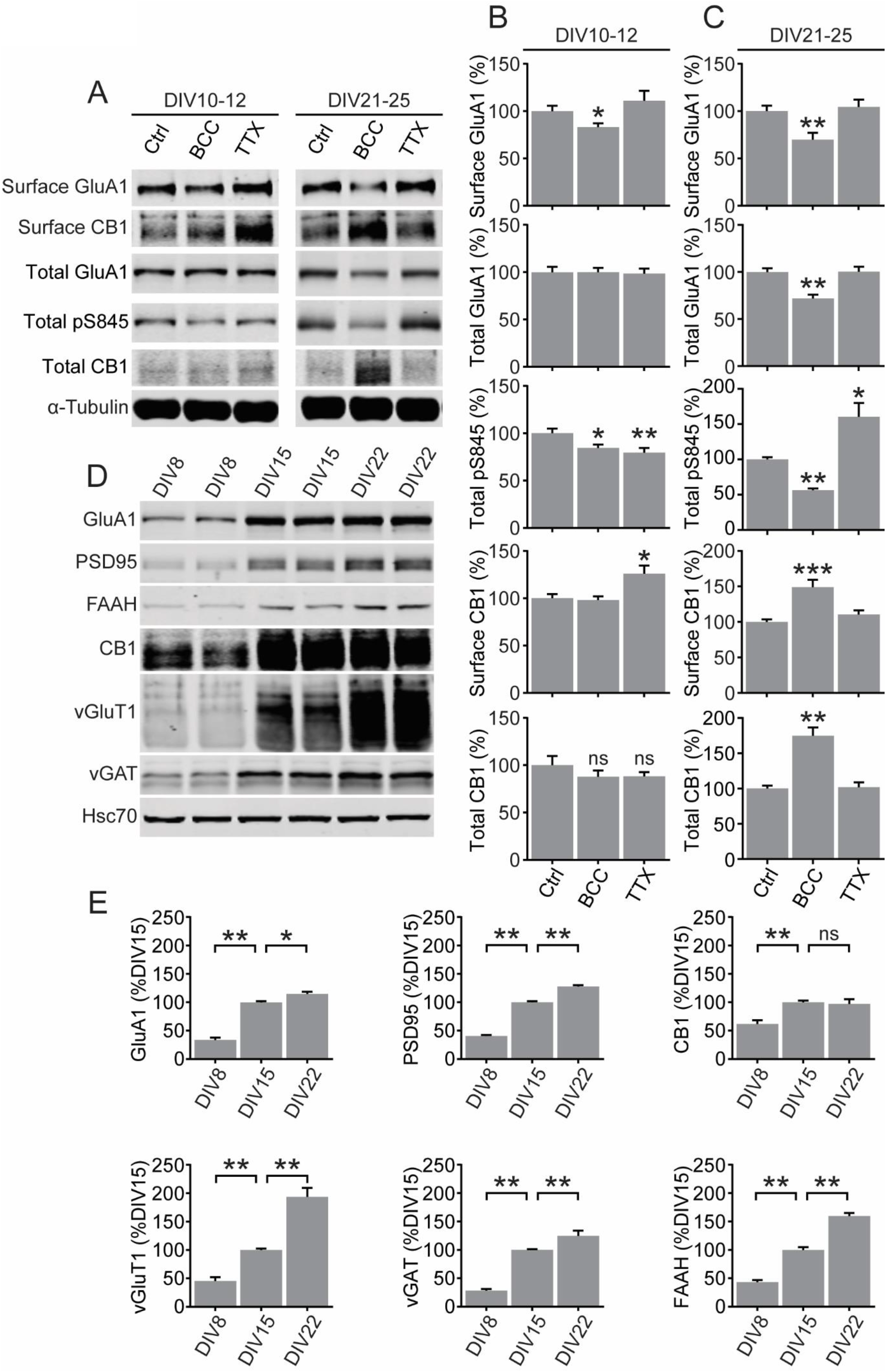
CB1 expression is upregulated during homeostatic downscaling in mature neurons. **(A)** Representative western blots of surface and total protein expression in developing (DIV10- 12) and mature (DIV21-25) cultured cortical neurons treated for 48h with control media (Ctrl), 20μM bicuculline (BCC) or 1μM tetrodotoxin (TTX). **(B,C)** Quantification of data shown in **(A)** in developing neurons **(B)** and mature neurons **(C)**. BCC induced downregulation of surface GluA1 and dephosphorylation of S845 regardless of age, but upregulation of total and surface CB1 only in mature neurons. Results were normalized for each protein to its expression under control levels and presented as mean ± SEM from at least three independent culture preparations with triplicate wells. **(D,E)** Representative western blots and quantification of total protein expression in cultured cortical neurons at DIV8, DIV15 and DIV22. Results are normalized for each protein to its expression levels at DIV15, and presented as mean ± SEM from at least three independent culture preparations with quadruplicate wells. Standard unpaired t-test was used for **(B,C)** and one-way ANOVA with Dunnett’s multiple comparison test was used for **(E)**. *p ≤ 0.05, **p ≤ 0.01

**Figure 2.**
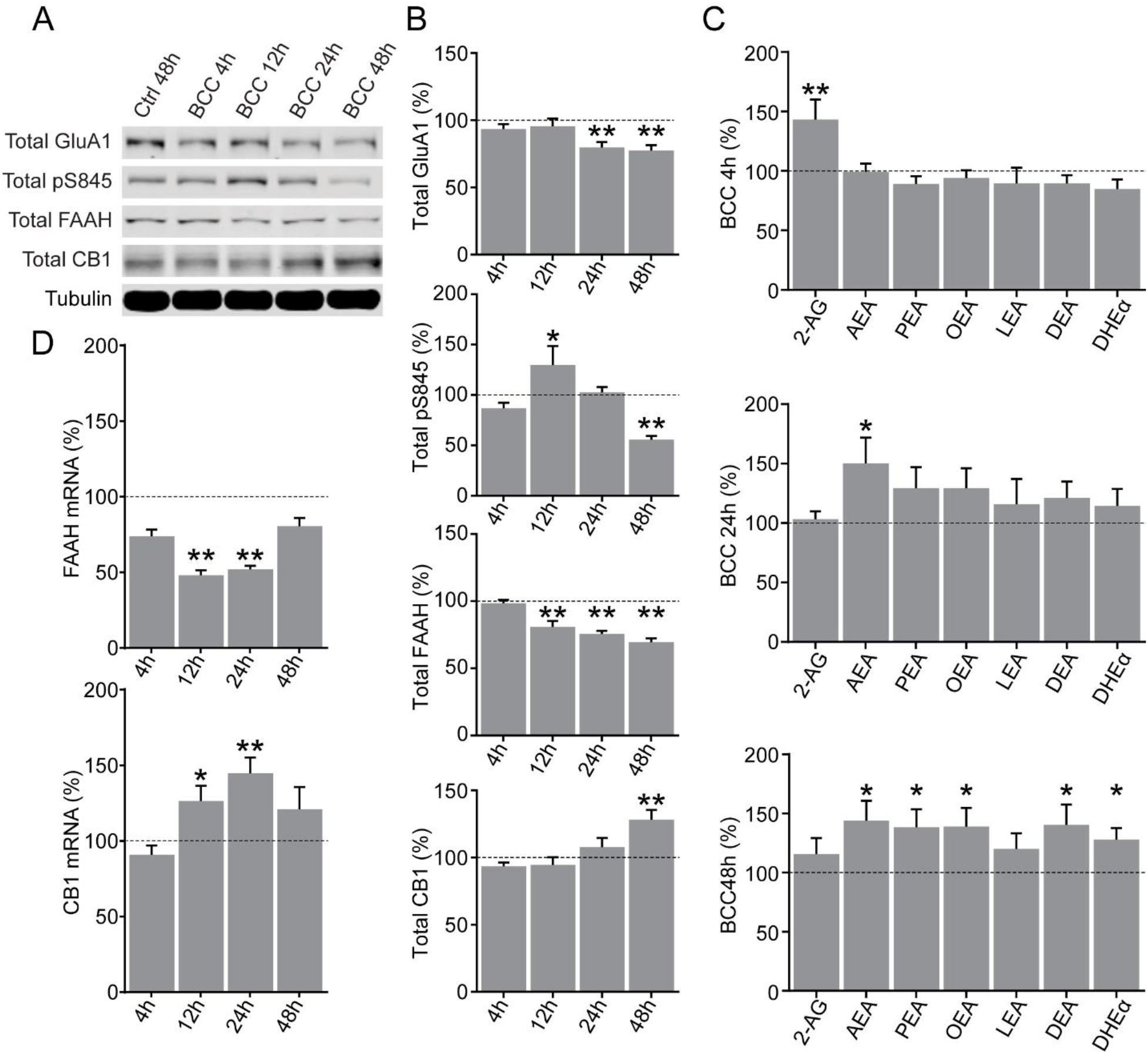
Time course of homeostatic downscaling in rat primary cortical neuron culture. **(A,B)** Representative western blots and quantification of total protein expression in DIV17 neurons treated by control media for 48h (Ctrl 48h) or by 20μM bicuculline (BCC) for 4h, 12h, 24h or 48h. FAAH is significantly downregulated from 12h and onward, but CB1 do not become significantly upregulated until the end of the 48h time course. Results are normalized for each protein to its expression levels in vehicle 48h (dotted line), and presented as mean ± SEM from four to six independent culture preparations with duplicate wells. **(C)** Quantification of targeted lipidomics analyzing 2-AG, AEA and various n-acylethanolamines (NAEs) in DIV17 neurons untreated or treated by 20μM BCC for 4h, 24h or 48h. Note that the neurons transition from a 2-AG dominated response at 4h to an AEA- and NAEs-dominated response at 24h and 48h, corresponding with the downregulation of FAAH shown in **(B)**. Results from each treatment group are normalized to a matching control, and presented as mean ± SEM from three to four independent culture preparations with duplicate plates. **(D)** Quantification of mRNA expression of FAAH and CB1 in DIV18-20 neurons untreated or treated by 20μM BCC for 4h, 24h or 48h. Results are normalized for each mRNA target to its expression levels in vehicle 48h (dotted line), and presented as mean ± SEM from three independent culture preparations with triplicate wells. One-way ANOVA with Dunnett’s multiple comparison test *p ≤ 0.05, **p ≤ 0.01

**Figure 3.**
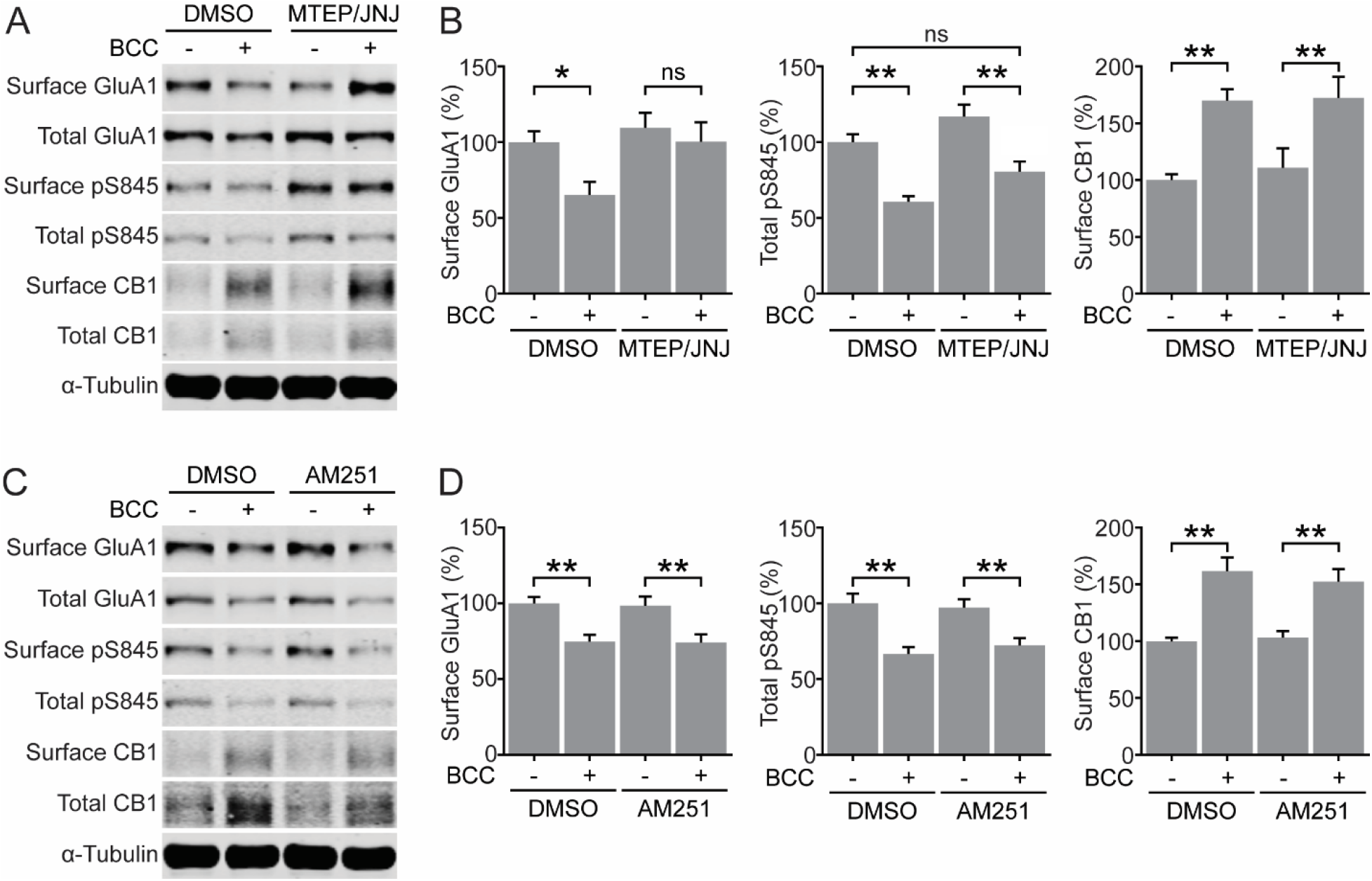
Glutamatergic and cannabinergic adaptations occur through independent mechanisms. **(A,B)** Representative blots and quantifications of surface and total protein expression in DIV21-25 neurons treated by DMSO or 20μM BCC in the absence or presence of 1μM MTEP hydrochloride, a selective mGluR5 antagonist, and 100nM JNJ 16259685, a selective mGluR1 antagonist (MTEP/JNJ). MTEP/JNJ blocked BCC-induced downregulation of surface GluA1 but not the upregulation of surface CB1. **(C,D)** Representative blots and quantifications of surface and total protein expression in DIV21-25 neurons treated by DMSO or 20μM bicuculline (BCC) in the absence or presence of 500nM AM251, a selective CB1 antagonist. AM251 did not block BCC-induced downregulation of surface GluA1 or the upregulation of surface CB1. Results are normalized for each protein to its expression levels in control group, and presented as mean ± SEM from four to six independent culture preparations with triplicate wells. One-way ANOVA with Šidák’s multiple comparison test *p ≤ 0.05, **p ≤ 0.01

**Figure 4.**
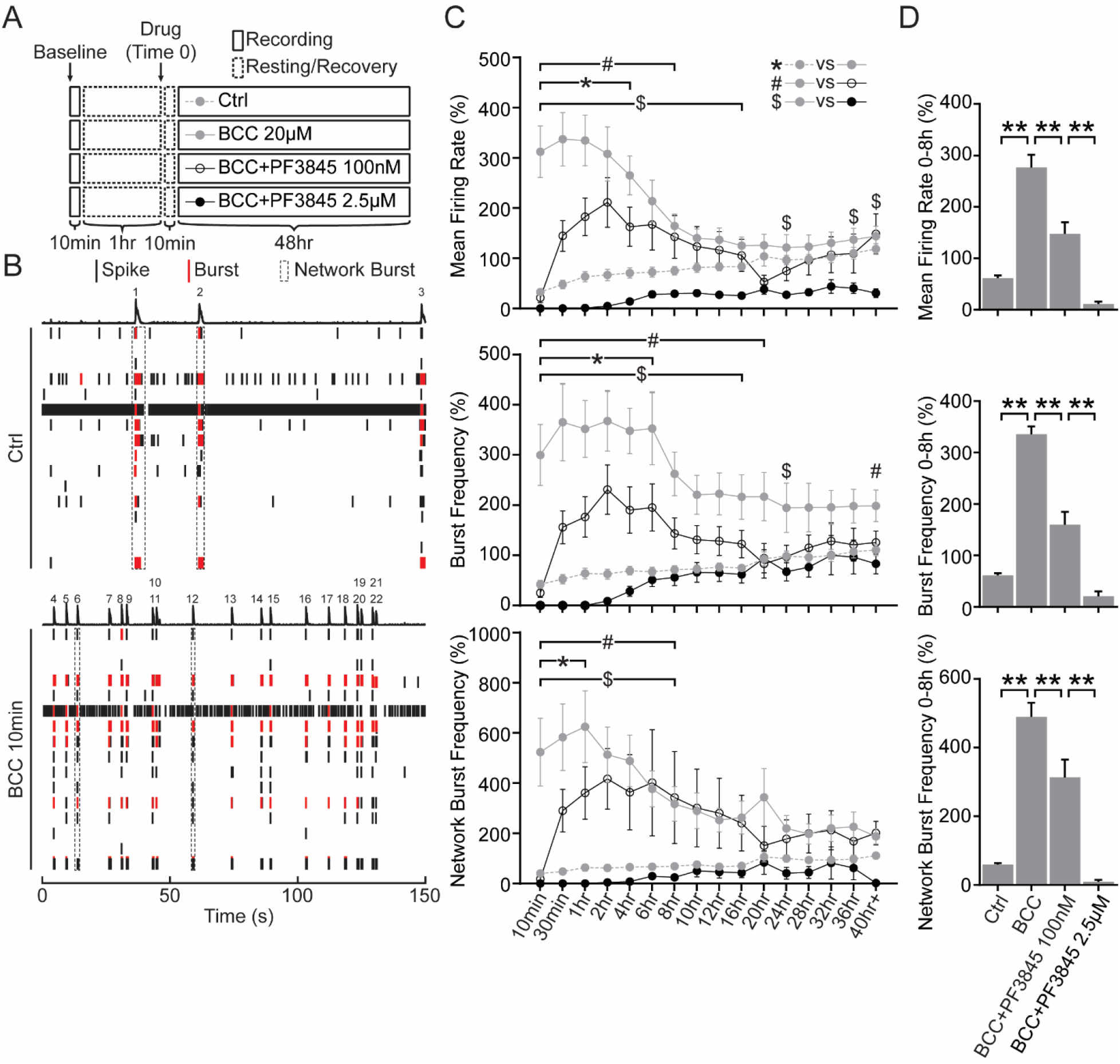
Elevated FAAH-substrates suppress network activities during homeostatic adaptation to BCC-induced hyperexcitation. **(A)** Experimental design. Neurons DIV21-28 were recorded for a 10min baseline, and then allowed to rest for 1hr. Network activities were then recorded for 10min once every 30min over 48h in neurons treated with ctrl media (Ctrl), 20μM BCC alone (BCC 20μM), or 20μM BCC in combination with PF3845, a potent FAAH inhibitor, at a lower dosage (BCC+PF3845 100nM) or a higher dosage (BCC+PF3845 2.5μM). Media was allowed to equilibrate for 10min after drug treatment (Time 0) before the first point was taken. **(B)** Raster plot of one well (16 electrodes) of MEA activities recorded for 150s at baseline (Ctrl) or 10min following 20μM BCC treatment (BCC 10min). Single electrode spikes (short black line) and single electrode bursts (short red line) are quantified as mean firing rate and burst frequency, while network bursts (dotted black box or numbered peaks) are quantified as network burst frequency, as shown in **(C)**. Note that 20μM BCC rapidly induces hyperexcitation in network activities within 10mins. This is exemplified here by 3 network bursts under control, but 19 after 10min of BCC. **(C)** Quantification of mean firing rate, burst frequency and network burst frequency, as illustrated in **(B)**, for neurons DIV21-28 treated for 48h with ctrl media, 20μM BCC, BCC+PF3845 100nM or BCC+PF3845 2.5μM, as illustrated in **(A)**. BCC treatment rapidly (<10min) converts the network to a burst firing pattern which remains significantly elevated for ∼8h, and the hyperexcitation can be suppressed by PF3845 in a dose-dependent manner. Results were normalized to a within-well control recorded for 10min at ∼1hr prior to treatment **(A**, baseline), and presented as mean ± SEM from six independent culture preparations with quadruplicate wells. Two-way ANOVA with Dunnett’s multiple comparison test ^#^p ≤ 0.05 Ctrl vs. BCC, *p ≤ 0.05 BCC vs. BCC+PF3845 100nM, ^$^p ≤ 0.05 BCC vs. BCC+PF3845 2.5μM throughout the duration indicated by the bracket and/or at time points denoted by the symbols. **(D)** Quantification of average mean firing rate, burst frequency and network burst frequency for the first 8h of treatment. Note that the dose-dependent suppression of network activities is particularly prominent in the first 8h during which the BCC-induced hyperexcitation remains statistically significant. One-way ANOVA with Tukey’s multiple comparison test *p ≤ 0.05, **p ≤ 0.01

**Figure 5.**
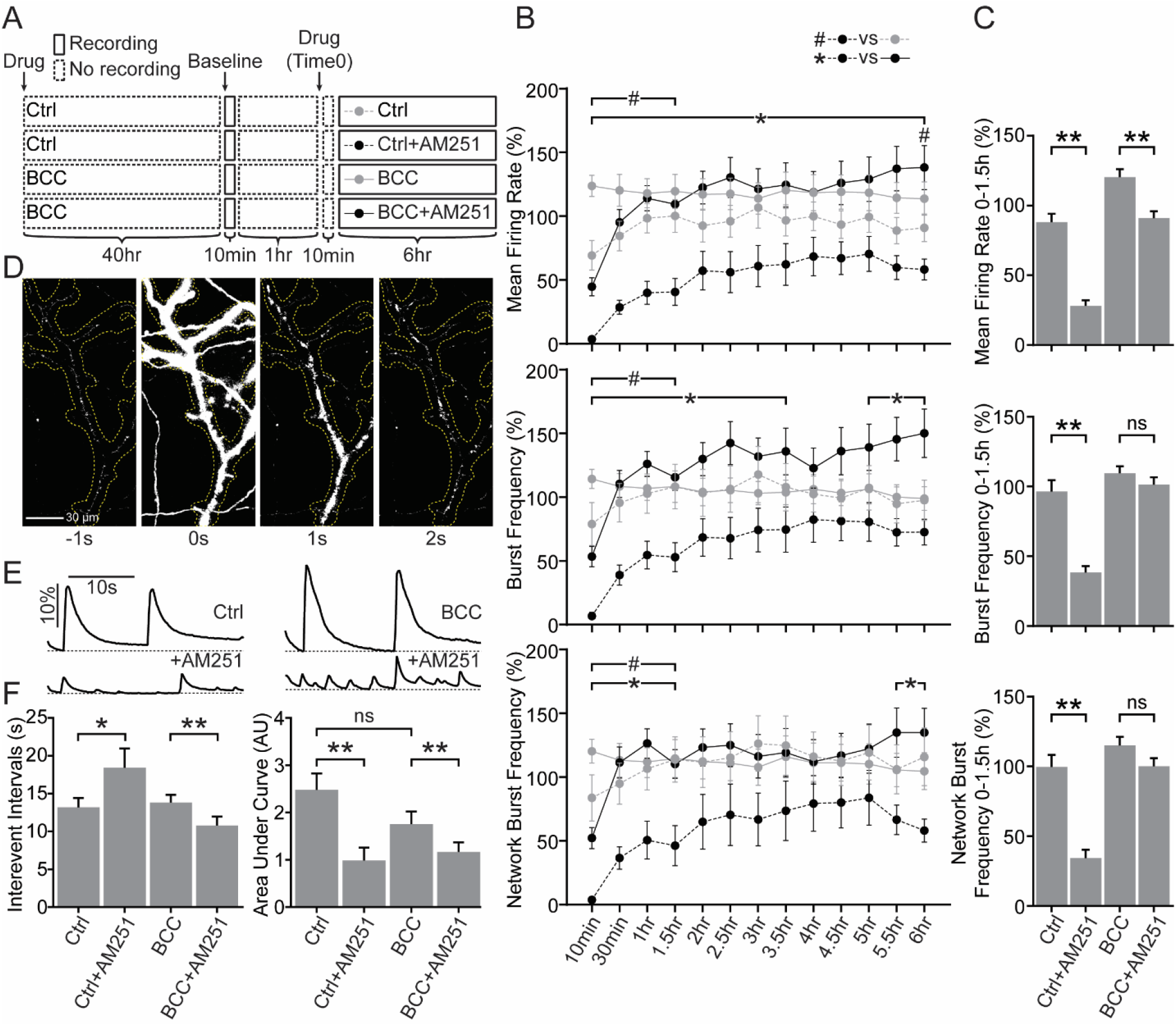
CB1 antagonism suppresses network activities before and after adaptation to BCC- induced hyperexcitation. **(A)** Experimental design. Neurons DIV21-28 were treated with ctrl media (Ctrl, or naive) or 20μM BCC (BCC, or downscaled) for 40hr before baseline was recorded for 10min. After resting for 1hr, naive or downscaled neurons were recorded and quantified with MEA in 10min bins once every 30min over 6hr in the absence (Ctrl, BCC) or presence (Ctrl+AM251, BCC+AM251) of 50nM AM251, a potent CB1 inverse agonist. Media was allowed to equilibrate for 10min after the drug treatment (Time 0) before the first time point was taken. The same experimental design was used for live-cell imaging, but those cells were only recorded for 1hr in the absence or presence of AM251. **(B)** Quantification of mean firing rate, burst frequency and network burst frequency from naive neurons untreated (Ctrl) or treated for 6h with 50nM AM251 (Ctrl+AM251), and from downscaled neurons untreated (BCC) or treated for 6h with 50nM AM251 (BCC+AM251), as illustrated in **(A)**. Note that AM251 strongly suppresses network activities in naïve cells which remains significantly depressed for 1.5h, but the response is greatly attenuated in downscaled cells. Results were normalized to a within-well control recorded for 10min at ∼1hr prior to treatment **(A**, baseline), and presented as mean ± SEM from four to six independent culture preparations with quadruplicate wells. Two-way ANOVA with Tukey’s multiple comparison test ^#^p ≤ 0.05 Ctrl vs. Ctrl+AM251, *p ≤ 0.05 Ctrl+AM251 vs. BCC+AM251 throughout the duration indicated by the bracket and/or at time points denoted by the symbols. **(C)** Quantification of average mean firing rate, burst frequency and network burst frequency for the first 1.5h of treatment, during which AM251-induced suppression of network activities remains statistically significant in naïve cells. Note that attenuation of AM251-induced suppression of network activities is particularly prominent during the first 1.5h. **(D)** Double derivative time-lapse images showing spontaneous changes in iGluSnFR fluorescence corresponding to a single synchronous glutamate event under control condition. **(E)** Representative traces of iGluSnFR fluorescence corresponding to multiple synchronous glutamate events recorded in DIV21-25 culture cortical neurons naïve and untreated (Ctrl, top left), naïve and treated with 500nM AM251 for up to 1hr (Ctrl+AM251, bottom left), downscaled and untreated (BCC, top right) or downscaled and treated with 500nM AM251 for up to 1hr (BCC+AM251, bottom right), similar to MEA experimental design illustrated in **(A)**. Results are presented as change in fluorescence for each recording normalized to its minimum fluorescence (ΔF/F, dotted line represent minimum fluorescence). Scale bar represents 10s (horizontal) and 10% change from minimum fluorescence (vertical). Ctrl data in panel is from the same set of data as in Extended Figure 5-1. **(F)** Quantification of interevent intervals and area under curve for data shown in **(E)**. Acute AM251 application causes reduction of glutamatergic event amplitude in naïve and downscaled cells, but leads to increased frequency in downscaled and reduced frequency in naïve cells. Results are presented as mean ± SEM from three to six independent culture preparations with four to eight wells in each culture. One-way ANOVA with Šidák’s multiple comparison test. *p ≤ 0.05, **p ≤ 0.01

### Antibodies

The following mouse primary antibodies were used: anti-GluA1 (NeuroMab 75-327, 1:1,000), anti-FAAH (Abcam 1:1,000), anti-PSD95 (NeuroMab 75-028, 1:1,000,000) and anti-α-Tubulin (Abcam, 1:50,000). The following rabbit primary antibodies were used: anti-GluA1 phospho- S845 specific (Millipore 1:1,000) and anti-CB1 (Cell Signaling, 1:1,000). The following guinea pig primary antibodies were used: anti-vGluT1 (Synaptic Systems 135 304, 1:1,1000), anti-vGAT (Synaptic Systems 131 005, 1:1,000). The following secondary antibodies were used: goat anti-mouse 680RD (Licor, 1:10,000), goat anti-rabbit 800CW (Licor, 1:15,000) and donkey anti-guinea pig 800CW (Licor, 1:15,000).

### Cell surface biotinylation

Surface biotinylation of cultured cortical neurons were performed as previously described [redacted for double-blind review]. Briefly, neurons were rinsed with ice-cold PBSCM (PBS containing 0.1mM CaCl_2_ and 1mM MgCl_2_, pH 8.0), incubated in PBSCM containing 0.5mg/mL Sulfo-NHS-SS-biotin (Thermo Scientific, 30min, 4℃), then rinsed in ice-cold PBSCM. Excess biotin was then quenched twice in PBSCM containing 20mM glycine (2 x 7min, 4℃), and then the cells were washed again with ice-cold PBSCM. Cells were lysed in ice-cold lysis buffer (PBS containing 1% Triton X-100, 0.5% sodium deoxycholate, 0.02% SDS, 1mM EDTA, 5mM sodium pyrophosphate, 1μM okadaic acid, 1mM Na_3_VO_4_ and phosphatase inhibitor cocktail [Roche]).

Lysates were pre-cleared with centrifugation (17,000G, 10min, 4℃). Protein concentration of each pre-cleared lysate was quantified using Bradford reagent (Bio-rad), and equal amounts of proteins were incubated overnight with NeutrAvidin-coupled agarose beads (Thermo Scientific). Beads were washed three times with ice-cold PBS containing 1% Triton X-100, and biotinylated proteins were eluted with 2x SDS sample buffer at 65℃ for 20min. Cell surface or total proteins were then subjected to SDS-PAGE and analyzed by western blot.

### Western blot

Lysates from cultured cortical neurons were pre-cleared with centrifugation (17,000G, 10min, 4℃). Protein concentration was determined using Bradford assay. Pre-cleared lysates were mixed with 2x SDS sample buffer (20% glycerol, 100mM Tris, 5% SDS, 5% BME, pH 6.8) and denatured at 65℃ for 20min. Equal amount of proteins were loaded and separated by SDS-PAGE on hand-cast gels. Separated proteins were transferred to a nitro-cellulose membrane (GE Healthcare). Following transfer, membranes were blocked in 3% BSA in TBS for 45min at room temperature. After blocking, membranes were incubated with primary antibodies dissolved in TBST (TBS containing 0.1% Tween-20) containing 3% BSA, overnight at 4℃. Membranes were then washed in TBST (3x 15min) before being incubated in secondary antibodies (Licor) dissolved in TBST containing 3% BSA and 0.01% SDS for 1 hr at room temperature.

Membranes were then washed again in TBST (3x 40min). Membranes were imaged on Licor Odyssey CLx Imaging System. Blots were analyzed and quantified using Image Studio software (Licor).

### Targeted mass spectrometry

Rat primary cortical neurons were grown on 10cm tissue culture plates at a density of 5,000,000 cells/plate for 17DIV. Culture media was aspirated, cells were rinsed with ice cold PBS and then collected and suspended by scraping in PBS containing 50nM JZL195 (Tocris), a dual inhibitor of FAAH and MAGL (Long et al., 2009). Cells were pelleted by centrifugation at 5,000xg for 5min, the supernatant was removed and cell pellets were flash frozen and stored at -80°C until further analysis. Frozen cell pellets were prepared for endocannabinoid analysis as follows.

Briefly, cell pellets were removed from -80°C freezer and thawed on ice. To each sample 170μl of methanol, 20μl of internal standard containing 200ng/ml each of arachidonyl ethanolamide-d4 and Oleoyl ethanolamide-d4 and 2000ng/ml of 2-arachidonlyl glycerol-d5, and 10μl of 5mg/ml BHT in ethanol was added. The cell pellet was resuspended and then vortexed for 5 seconds.

The sample was then centrifuged at 14,000RPM for 10 minutes at 4°C. The supernatant was removed and then placed into a capped autosampler vial for analysis. LC/MS/MS analysis of endocannabinoids was performed as previously described (Gouveia-Figueira and Nording, 2015), with some modifications. Briefly, mass spectrometric analysis was performed on an Agilent 6490 triple quadrupole mass spectrometer in positive ionization mode. Calibration standards were analyzed over a range of concentrations from 0.025–50pg on column for all of the ethanolamides and 2.5-5000pg on column for the 2-AG. The following lipids were quantified: 2-arachidonyl glycerol (2-AG), arachidonoyl ethanolamide (AEA), docosahexaenoyl ethanolamide (DHEa), docosatetraenoyl ethanolamide (DEA), linoleoyl ethanolamide (LEA), oleoyl ethanolamide (OEA), palmitoleoyl ethanolamide (POEA), palmitoyl ethanolamide (PEA), stearoyl ethanolamide (SEA). Quantitation of endocannabinoids was performed using Agilent Masshunter Quantitative Analysis software. All results were normalized to protein concentration.

### Quantitative real-time PCR

mRNA was purified using the RNeasy Plus Mini Kit (Qiagen) according to the manufacturer’s instructions. RNA was reverse transcribed into cDNA using the High-Capacity cDNA Reverse Transcription kit (Applied Biosystems). Quantitative real-time PCR (RT-qPCR) was performed using the QuantStudio 7 Flex (Applied Biosystems) using TaqMan Fast Advanced master mix and TaqMan primer/probes (Invitrogen) for GAPDH (Rn01775763_g1), FAAH (Rn00577086_m1), CB1 (Rn02758689_s1). Expression data was calculated as fold gene expression using 2^−ΔΔCt^ method with GAPDH as the reference gene, and normalized to untreated control. Four independent technical replicates were performed in each individual experiment.

### Multi-electrode Array

Dissociated rat cortical neurons were prepared as above and plated at 120,000 cells/well onto 24-well Cytoview plates (Axion Biosystems) coated with 0.1 mg/mL PLL (Sigma-Aldrich). The plating area fully covered the bottom of the well. Recordings were performed. Neurons were maintained as above until DIV21 when they were treated with drugs and used for experiments. Multielectrode array (MEA) recordings were performed in culture media using a Maestro Edge system and AxIS software (Axion Biosystem; version 3.2), with a bandwidth filter from 200 Hz to 3.0 kHz. Spike detection was computed with an adaptive threshold of 6 times the standard deviation of the estimated noise for each electrode. Plates were left untouched in the Maestro instrument for 10min prior to recording, which proceeded for 10min once every 30min for as long as needed in each experiment. Data were analyzed using the neural metrics tool (Axion Biosystems; version 3.1.7), under the conditions that an electrode was deemed active if 5 or more spikes occurred over 1min. The mean firing rate for each well was defined as number of spikes divided by the duration of the analysis, in Hz. Single electrode bursts were defined as bursts of a minimum of 5 spikes on one electrode with a maximum inter-spike interval (ISI) of 100ms, which was divided by the duration of the analysis to yield burst frequency, in Hz. Network bursts were defined as bursts of a minimum of 35 spikes that occurred in more than 35% of the active electrodes in the well, with a maximum ISI of 100ms, which was divided by the duration of the analysis to yield network burst frequency, in Hz.

### Transfection and Live-cell wide field microscopy

A mixture of 1.5 μL of Lipofectamine 2000 (Invitrogen) and 2.0 μg of pAAV.CAG.SF-iGluSnFR.A184S (Addgene plasmid # 106198 ; gifted from Loren Looger; http://n2t.net/addgene:106198 ; RRID:Addgene_106198) in 60 μL Neurobasal Medium (Gibco) was prepared and incubated at room temperature for 30 minutes. Media from neuron cultures was saved aside and cells were incubated with this mixture at 37° C and 5% CO_2_ for 30 minutes. Mixture was aspirated and replaced with original media. Time lapse wide field fluorescence videos were obtained at 5 frames per second for 60s using a Zeiss Laser Scanning Microscope (LSM) 800/Airyscan equipped with a Colibri 7 LED light source, Zeiss Axiocam 506 CCD camera, and 63x/1.4 NA objective lens at 37° C and 5% CO_2_. Neurons were imaged in their culture media.

### Live-cell Imaging Data Analysis

Timelapse images were analyzed in FIJI by first subtracting background fluorescence measured from an empty region of the field of view then running an exponential fit bleach correction using a region of interest containing dendrites of a cell in the field of view. The dendritic arbor was traced in FIJI and designated the region of interest. Fluorescence levels in this region of interest were measured and analyzed semi-automatically with a custom code in R, which identifies frames where synchronized fluorescent changes began by locating peaks in the second derivative of a plot of fluorescence over time. A baseline fluorescence was assigned to each synchronous event, corresponding with the fluorescence intensity value of the first frame preceding the synchronous event and change in fluorescence over background (ΔF/F) over baseline was calculated for each frame. The area under the curve of synchronous events was measured in arbitrary units, by summing the ΔF/F values for each frame from the initial rise from baseline to the return to baseline. Inter-event interval was calculated by counting the number of frames from the start of one event to the start of the next and converting this value to seconds based on recording frame rate. Statistics were run in Prism. For presentation purposes only, a double derivative time lapse was generated that allows for the clear visualization the change in fluorescence with time.

### Study design and statistical analysis

Results are presented as mean ± SEM from at least three independent culture preparations. Statistical significances were determined using unpaired t tests between Ctrl, BCC and TTX. One-way ANOVA and appropriate *post hoc* multiple comparisons were used, as indicated in figure legends, for comparisons of mRNA, protein and targeted lipidomics at different time points in downscaling time course, for comparisons between Ctrl and BCC with or without MTEP/JNJ or AM251, for comparisons of iGluSnFR interevent intervals and area under curve before and after acute AM251 treatment, for comparisons of average network activities for the first 8h between Ctrl, BCC and BCC+PF3845 treated cells, and for the comparisons of average network activities for the first 1.5h between naïve and downscaled cells treated with AM251.

Two-way ANOVA and appropriate *post hoc* multiple comparisons were used, as indicated in figure legends, for comparisons of network activities between cells treated with Ctrl, BCC and BCC+PF3845, and for comparisons of network activities between naïve and downscaled cells treated with AM251. All statistical analyses were performed in GraphPad Prism.

## Results

### Adaptation of the eCB system during homeostatic downscaling

The endocannabinoid system (eCB) is a major neuromodulatory system that regulates neurotransmitter release at both glutamatergic and GABAergic synapses. To determine a possible role of the eCB system in homeostatic plasticity, developing (DIV11-12) and mature (DIV21-25) cultured rat cortical neurons were treated with bicuculline (BCC) or tetrodotoxin (TTX) for 48hrs, treatments previously shown to robustly induce homeostatic downscaling and upscaling respectively (Turrigiano et al., 1998). Adjustments in surface AMPAR receptor density and the phosphorylation status of AMPAR subunit GluA1 at S845 are previously described markers for synaptic scaling (Diering et al., 2014; Hu et al., 2010; Kim and Ziff, 2014; O’Brien et al., 1998). Neurons were surface biotinylated followed by lysis to allow for the detection of total and surface proteins (Fig. 1A). We examined the expression of AMPAR receptor subunit GluA1 and cannabinoid receptor 1 (CB1). In developing and mature cultures, BCC induced a significant downregulation in surface GluA1 and dephosphorylation of GluA1 S845 (pS845), indicating that the neurons have engaged homeostatic downscaling, consistent with previous findings (Fig. 1A-C). Network suppression with TTX caused a trend of increased surface GluA1 as expected, although these changes were not significant, and in mature cultures we observed the expected TTX-induced increase in phosphorylation of GluA1 S845 previously associated with homeostatic upscaling (Diering et al., 2014; Kim and Ziff, 2014). In developing cultures BCC had no effect on CB1 expression, whereas TTX treatment caused a significant increase in surface, but not total, CB1 levels (Fig. 1B). In contrast, mature cultures responded to BCC by a significant upregulation of surface and total CB1 expression, compared to control, whereas TTX had no effect (Fig. 1C). The above data suggest that eCB system adapts to changes in network activity in a manner that depends on the maturation of the culture.

To further investigate the maturational changes in vitro, we measured synapse and endocannabinoid system protein expression in neurons at DIV8, DIV15 and DIV22 using western blot (Fig. 1D). In the second week in vitro (DIV8 vs. DIV15), we found a dramatic increase in all proteins examined (Fig. 1E). In the third week in vitro (DIV22 vs. DIV15), we found a striking increase in the expression of vesicular glutamate transporter 1 (vGluT1) and fatty acid amid hydrolase (FAAH), the principal degradative lipase for anandamide (AEA) and related N-acylethanolamines (NAEs). Additionally, we found a moderate increase in GluA1, PSD95, and vesicular GABA transporter (vGAT) and no change in CB1 (Fig. 1E). These results suggest that, compared to developing neurons, mature neurons further undergo a significant amount of synapse maturation, especially in pre-synaptic glutamate release machinery such as vGluT1, and potentially have significantly different metabolism of NAEs as indicated by increased expression of FAAH. In our subsequent experiments we have focused on the adaptation of the eCB system to network hyperexcitability in mature neuron cultures.

### Temporal Dynamics of eCB Adaptations

To further characterize the adaptations in the eCB system during homeostatic downscaling in mature cultures, we performed a time course of BCC treatment (4-48hrs) followed by RT-qPCR and western blot analysis of transcript and protein expression respectively, and quantification of eCB lipids using targeted mass spectrometry. Total GluA1 showed a significant decrease beginning at 24hrs as expected. pS845 increased over the first 12 hours but decreased significantly at 48h (Fig. 2A and B). CB1 protein increased significantly but only at 48hr, suggesting that this was a delayed response (Fig. 2B). Interestingly, FAAH protein decreased progressively starting at 12hr and remained significantly downregulated through the 48hr time course, suggesting that adaptations in eCB metabolism are an earlier step in homeostatic downscaling (Fig. 2B). Accordingly, targeted mass spectrometry of eCBs showed that while 2-AG upregulation was a prominent feature of BCC treatment at 4hr, a significant upregulation of AEA became the dominant response at 24hr and onward. By 48hr, multiple NAEs, all of which are FAAH substrates, were significantly upregulated (Fig. 2C). The BCC-induced decrease in FAAH protein was matched by a decrease in the expression of FAAH mRNA at 12-24hrs, suggesting that changes in FAAH protein are controlled in part at the level of transcription. Interestingly, BCC treatment induced an upregulation of CB1 mRNA at 12-24hrs post BCC treatment, well in advance of CB1 protein increases at 48hrs, suggesting that CB1 protein upregulation is controlled by both transcriptional and translational mechanisms in response to hyperexcitation. Together, these findings suggest that during the induction and expression of homeostatic downscaling in mature cortical cultures, there is a stepwise adaptation in the eCB system starting with downregulation of FAAH protein and consequent increases in the abundance of AEA and other NAEs, followed by the delayed upregulation of CB1. Changes in AEA metabolism and upregulation of CB1 likely synergize to enhance tonic eCB signaling.

### Coordination of presynaptic CB1 signaling and postsynaptic glutamatergic signaling

Homeostatic downscaling in cultured rodent neurons is primarily understood as a post- synaptic adaptation that involves downregulation of synaptic AMPARs. Since BCC-induced up- regulation of CB1 receptors suggests a coordinated pre- and post-synaptic response, we next performed a series of pharmacology experiments to examine the relationship between post- synaptic glutamatergic signaling and pre-synaptic cannabinergic signaling during BCC-induced downscaling. A previous report has shown that hyperexcitation caused by BCC resulted in activation of type I mGluR1/5 and that a cocktail of non-competitive mGluR1/5 inhibitors could prevent post-synaptic downscaling of AMPARs (Hu et al., 2010). We confirmed this finding by treating neurons with a cocktail of 1μM MTEP and 100nM JNJ16259685 (MTEP/JNJ), non-competitive inhibitors of mGluR5 and mGluR1 respectively, and showing that this treatment prevented BCC-induced downregulation of surface GluA1 and partially blocked the dephosphorylation of pS845 (Fig. 3B). However, MTEP/JNJ treatment had no effect on BCC- induced upregulation of total and surface CB1 (Fig. 3B). This result suggests that pre-synaptic adaptation of CB1 during downscaling is independent of post-synaptic glutamatergic signaling.

Next, we blocked cannabinergic signaling in cultured rat cortical neurons by treating with selective CB1 antagonist AM251 (500nM) and examined the effect on post-synaptic adaptation during downscaling. AM251 treatment had no effect on surface GluA1 and pS845 and did not prevent the BCC-induced downregulation of surface GluA1 and pS845 (Fig. 3D). Interestingly, the BCC-induced upregulation of CB1 was not prevented by AM251, suggesting that CB1 upregulation is also independent of CB1 signaling. Together, these findings suggest that neurons respond to network hyperexcitability with coordinated cannabinergic and glutamatergic adaptations, which occur through independent molecular mechanisms.

### Enhancing eCB signaling suppresses network activity during hyperexcitation

In the context of network activity, CB1 signaling is well characterized to regulate pre- synaptic vesicle release dynamics (Castillo et al., 2012), and to maintain excitatory up-states, an intrinsic mode of cortical network activity particularly prominent in sleeping and anesthetized animals (Pava et al., 2014; Steriade et al., 1993a). Since up-state like synchronous activity is also well documented in organotypic slices (Johnson and Buonomano, 2007; Sanchez-Vives and McCormick, 2000) and cultured cortical neurons (Canepari et al., 1997; Kamioka et al., 1996; Saberi-Moghadam et al., 2018), the reorganization in eCB system described above (Fig. 2A-D) may be involved in the regulation of network activity. To better understand the role of eCBs in network activity, we monitored spiking activities from neurons plated on multi-electrode arrays (MEA) as neural networks adapt to BCC-induced hyperexcitation (Fig. 4A). Mean firing rate was quantified as total number of spikes divided by the duration of the analysis. Single electrode bursts (Fig. 4B, red line) and networks bursts (Fig. 4B, dotted black box) were identified based on spike densities and participating electrodes (see methods), the total of which was then divided by the duration of the analysis to yield burst frequency and network burst frequency, respectively.

In agreement with previous studies, we observed that mature neurons (>DIV21) exhibited baseline levels of spontaneous network activation characterized by highly rhythmic windows of coordinated firing (“network bursts”) followed by periods of relative network silence (Fig. 4B, control condition). BCC application rapidly (<10min) converted the network activity to a burst firing pattern (Fig. 4B and C). The mean firing rate and network burst frequency peaked around 1-2hr and remained significantly elevated for ∼8hr, after which they became statistically indistinguishable from the baseline, suggesting that some homeostatic adjustments had occurred within 8h of BCC treatment (Fig. 4C, top, bottom, ^#^p<0.05). The single-electrode burst frequency also peaked around 1-2hr but remained significantly elevated for 20hr (Fig. 4C, middle, ^#^p<0.05). The recovery of network activity within 8h of hyperexcitation onset likely reflects mechanisms of homeostatic processes other than downregulation of synaptic AMPARs and excitatory drive which is usually expressed after chronic (>24h) perturbations in network activity (Fig. 2B) (Diering et al., 2014; Fong et al., 2015; Gonzalez-Islas et al., 2020).

Since decreases in AEA degradation could be detected early on (Fig. 2B), we hypothesized that the downregulation of FAAH and upregulation of AEA may be involved in network adaptation to BCC-induced hyperactivity. Accordingly, increasing AEA/NAE availability by application of selective FAAH inhibitor PF3845 (Ahn et al., 2009) suppressed BCC-induced network hyperexcitation in a dose-dependent manner, resulting in lowered mean firing rate, burst frequency and network burst frequency comparing to BCC only treatment (Fig. 4C, *p<0.05, ^$^p<0.05). The dose-dependent suppression of hyperexcitation is particularly prominent during the first 8hr of BCC treatment (Fig. 4D). This result suggests that downregulation of FAAH, and upregulation of AEA, is part of a mechanism to reduce network excitability in response to BCC-induced hyperexcitation.

### Inhibiting CB1 signaling and excitatory network activity

Since the eCB system is involved in the regulation of network activity (Fig. 4C and D), we hypothesized that the delayed upregulation of CB1 after BCC treatment described above (Fig. 1C and Fig. 2B) predicts a different network response to CB1 antagonism before and after 48h BCC treatment. To examine the role of CB1 on network activity following chronic BCC treatment, we left neurons untreated (naïve) or treated with BCC (downscaled) for 40hrs, and then monitored changes in spiking activities during 6h of CB1 inhibition using 50nM AM251 (Fig. 5A). In naïve cells, CB1 antagonism consistently led to a transient but significant decrease in all activity metrics examined for the first 1.5h, indicating acute suppression of network activities (Fig. 5B, ^#^p<0.05). However, in downscaled cells, AM251-induced acute suppression of network activities was negligible and virtually undetectable (Fig. 5B), except a minor decrease in mean firing rate throughout the first 1.5h (Fig. 5C, top). A direct comparison between control and BCC- treated cells showed a robust difference in mean firing rate and burst frequency responses to AM251 throughout 6h (Fig. 5B, *p<0.05), and a drastic difference in burst frequency and network burst frequency during the first 1.5h (Fig. 5C).

Since CB1 is known to regulate the release of neurotransmitters including glutamate, we examined whether changes in glutamate release were involved in network adaptations. To this end, we indirectly measured excitatory network activities by visualizing glutamate release dynamics with a high-affinity variant of fluorescent glutamate biosensor iGluSnFR-A184S (Marvin et al., 2013; Marvin et al., 2018). iGluSnFR is a plasma-membrane-targeted fluorescent reporter that responds to extracellular glutamate release with an increase in fluorescence (Marvin et al., 2018). We expressed iGluSnFR in mature neurons (>DIV21) using lipofection and examined glutamate release dynamics in live-cells using wide-field microscopy time-lapse recording. Glutamate release is quantified as a transient increase in iGluSnFR fluorescence (ΔF/F).

Under control conditions, we observed spontaneous synchronous events where fluorescence across the entire dendritic arbor was increased (Fig. 5D). The “wave-like” synchronous events were highly rhythmic, reminiscent of network burst activity recorded by MEA. Indeed, synchronous iGluSnFR events were completely suppressed by the sodium channel blocker TTX and reduced in amplitude by the NMDAR inhibitor D-AP5 (Ext. Fig 5-1) matching previously reported effects of these drugs on MEA network events (Sanchez-Vives and McCormick, 2000; Steriade et al., 1993b). Therefore, we suggest that synchronous iGluSnFR events are analogous to synchronous network burst events, or “cortical up-states” recorded by MEA. Moreover, acute BCC treatment (∼10min) resulted in a plateau in iGluSnFR synchronous events, suggesting a transiently sustained glutamate release indicative of hyperexcitation (Ext. Fig. 5-1), consistent with the transient increase in bursting activity measured by MEA recordings (Fig. 4B and C). Following 48h chronic BCC treatment, the plateau of synchronous events was absent, and the frequency and amplitude of synchronous events were comparable to control level, indicative of homeostatic adjustment of synchronous glutamate release (Fig. 5E and F). The similar pharmacological responses and homeostatic adjustment likely reflect that the synchronous glutamatergic events are, to a certain degree, analogous to up-state network events detected by MEA.

To examine the effect of CB1 antagonism on glutamate release following the induction of downscaling, we acutely blocked CB1 with AM251 for 1hr in naïve, or BCC-treated neurons (Fig. 5C). In both naive and downscaled cells, acute AM251 significantly suppressed the magnitude of synchronous glutamate release events (Fig. 5E), as measured by decreased area under the curve (Fig. 5F, right). Further, in naïve cells CB1 inhibition reduced the frequency of glutamatergic events, whereas in downscaled cells, AM251 application caused glutamatergic events of low amplitude to become more frequent (Fig. 5E and F, left). Together, these data suggest that CB1 inhibition has a diminished capacity to suppress network events after downscaling.

## Discussion

The eCB system has been well described to participate in multiple forms of short- and long-term synaptic plasticity (Castillo et al., 2012). However, to our knowledge, homeostatic adaptations of the eCB system in vitro have only been studied in the context of chronic network silence (Kim and Alger, 2010; Song et al., 2015). In this study we provide quantitative analysis of the adaptations of the eCB system to network hyperexcitability induced by chronic BCC treatment in cultured rat cortical neurons, an *in vitro* system that has been used to investigate the molecular mechanisms of homeostatic downscaling (Diering et al., 2014; Hu et al., 2010; Turrigiano et al., 1998). We report evidence of rapid and persistent downregulation of FAAH, consequent upregulation of bioactive NAEs including AEA, and delayed upregulation of CB1.

Furthermore, we show that pre-synaptic cannabinergic adaptations and post-synaptic glutamatergic adaptations occur through independent mechanisms. Finally, we show that endocannabinoid signaling through CB1 stabilizes network activities during homeostatic adaptation to hyperexcitation (Fig. 6).

**Figure 6.**
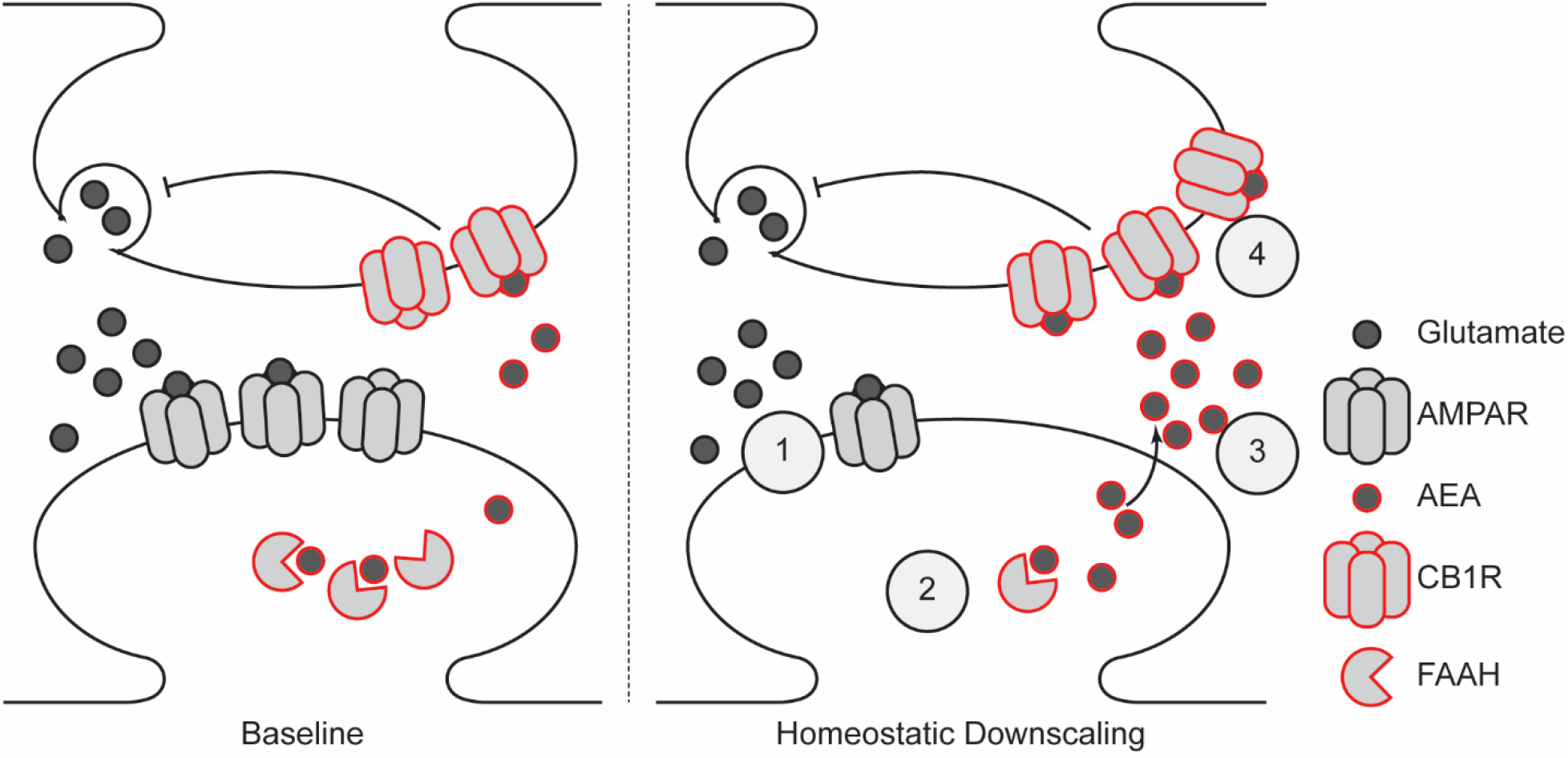
Model of homeostatic adaptation of the endocannabinoid system during downscaling. BCC induced downscaling drives coordinated remodeling of synapses: 1. Down regulation of surface AMPARs, 2. Down regulation of FAAH, 3. Accumulation of FAAH substrates including AEA, and 4. Up-regulation of surface CB1 receptors. Elevated FAAH substrates may serve to suppress network during adaptation to hyperexcitation, but CB1 signaling is also required to maintain network activities at baseline.

In developing cultures ∼2 weeks in vitro, homeostatic plasticity predominantly reflects changes in post-synaptic receptor density (Diering et al., 2014; Turrigiano, 2008). Additional pre-synaptic homeostatic adaptations are engaged as neurons mature in culture ∼3 weeks in vitro (Wierenga et al., 2006). Similarly, our data show that the more mature 3-week-old neuron cultures respond to chronic hyperexcitation in part through upregulation of surface and total CB1 (Fig.1). Other adaptations in mature cultures include homeostatic regulation of vesicular transporters (De Gois et al., 2005). Compared to developing cultures, mature cultures undergo further synaptic, dendritic, and functional development, which may confer greater need to regulate pre-synaptic components and network activities during homeostatic downscaling

(Harrill et al., 2015; Wagenaar et al., 2006). Mature cultures show more developed synapses as evidenced by increased expression of post-synaptic ion channel GluA1 and pre-synaptic vesicle release machineries vGluT1 and vGAT (Fig.1E). Although total CB1 levels were similar at DIV15 and DIV21, we observed a clear increase in the expression of FAAH, indicating that the eCB system is still undergoing maturation. We suggest that the upregulation of FAAH, and the emergence of the BCC-induced upregulation of CB1 during the maturation of cortical neurons in vitro, may reflect the developmental emergence of the need for activity-dependent regulation of synchronous network activities (Johnson and Buonomano, 2007; Kamioka et al., 1996).

### 2-AG versus AEA

2-AG and AEA are both well described agonists of the CB1 receptor. However, these two metabolites have completely non-overlapping enzymes for synthesis and degradation, suggesting that the two most prominent eCBs undergo distinct regulation and serve separate physiological functions (Di Marzo, 2018). 2-AG mediates major forms of eCB-dependent plasticity, such as DSI, DSE and mGluR-dependent LTD (Castillo et al., 2012; Tanimura et al., 2010; Yoshino et al., 2011). However, AEA has been implicated in cortical up-sate activity, sleep stability and memory (Busquets-Garcia et al., 2011; Murillo-Rodriguez et al., 1998; Pava et al., 2014), and AEA but not 2-AG are increased in cortex during the sleep phase (Martin et al., 2022). Our results indicate that AEA but not 2-AG exerts tonic action during homeostatic downscaling (Fig. 2). Some properties make AEA more suitable for tonic signaling. First, AEA is considered a partial agonist at CB1, which makes it less efficacious than full agonists such as 2- AG (Burkey et al., 1997; Glass and Northup, 1999; Sugiura et al., 1996). By regulating AEA during global downscaling of synaptic strength, neurons may preserve sensitivity to 2-AG-mediated “fine-tuning” of individual synapses. Second, sustained 2-AG signaling results in desensitization of CB1, whereas sustained AEA signaling does not (Schlosburg et al., 2010). By minimizing receptor desensitization, neurons may maintain tonic endocannabinoid signaling throughout the relatively long time-frame of homeostatic plasticity and avoid functional antagonism due to sustained 2-AG signaling, while at the same time preserving the short-term plasticity mediated by 2-AG. Indeed, Kim and Alger (2010) showed that chronic inactivity in hippocampal slices reduces endocannabinoid tone by regulating AEA degradation instead of 2- AG. Therefore, we hypothesize that 2-AG and AEA indeed mediate distinct mechanisms: a phasic action of 2-AG mediating short term plasticity, and a tonic action of AEA that regulates neuronal network activity over longer time scales.

Our time course data suggest that multiple FAAH substrates have become significantly upregulated by 48h (Fig. 2C). Since FAAH is the major enzyme for catabolizing multiple endocannabinoid substrates in the NAE family, it is possible that cellular levels of NAEs are regulated through a common mechanism to mediate coordinated signaling events (Di Marzo, 2018). Two such NAEs, oleoylethanolamide (OEA) and palmitoylethanolamide (PEA), are agonists for nuclear endocannabinoid receptor PPARα, and have been implicated in sleep-wake regulation and memory acquisition (Mazzola et al., 2009; Murillo-Rodriguez et al., 2011).

Indeed, PPARα has been described to be required for expression of the immediate early gene Arc in response to hippocampal neuronal activity through CREB-mediated transcriptional control of plasticity-related molecules (Roy et al., 2013). Thus, we speculate that AEA-CB1 signaling is only one part of a coordinated neuronal response that synergizes with signaling pathways activated by other bioactive NAEs. A possible role for OEA, PEA and/or PPARα should be investigated in future studies.

### CB1, cortical up-states and NREM sleep

In the absence of external stimuli, such as during anesthetization, NREM sleep or quiet wake, the mammalian neocortex shows intrinsic network oscillation between periods of synchronized depolarization (up-state) and prolonged network silence (down-state) (Steriade et al., 1993b; Timofeev et al., 2001). While we do not presume that dissociated cortical culture is identical to rodent neocortex in every aspect, cortical up-states and network activities share similar characteristics across a wide-range of *in vivo* or *in vitro* preparations (Johnson and Buonomano, 2007; Kaufman et al., 2014; Pava et al., 2014; Sanchez-Vives and McCormick, 2000; Steriade et al., 1993b; Timofeev et al., 2001). The similarities in these features across studies suggest that oscillatory activities may be a fundamental property of cortical networks. The common mechanisms may be studied in more convenient reduced preparations, such as dissociated cortical cultures, where greater experimental control can be exerted. In this context, our work adds to a growing body of literature that explores downscaling mechanisms (Diering et al., 2014; Diering et al., 2017) and/or network dynamics (Hinard et al., 2012; Kaufman et al., 2014; Mikhail et al., 2017; Saberi-Moghadam et al., 2018) in sleep-wake regulation using in vitro preparations. Given the ubiquity of eCB signaling in mammalian neocortex (Busquets-Garcia et al., 2018; Di Marzo, 2018), its diurnal variation (Murillo-Rodriguez et al., 2006; Valenti et al., 2004) and its known involvement in sleep-wake regulation (Martin et al., 2022; Pava et al., 2016), it is of interest to understand the mechanisms through which eCB signaling may influence prominent sleep features such as cortical up-states and network oscillations.

In the present study, we report that up-state like network activity in dissociated cortical neuron culture responds to BCC-induced hyperexcitation, followed by an eventual return to baseline, indicative of homeostatic regulation (Fig. 4C). Similar return to synchrony and homeostatic adjustments of network activities have been reported (Bateup et al., 2013; Kaufman et al., 2014; Saberi-Moghadam et al., 2018). Here we show a dose-dependent suppression of BCC-induced network hyperexcitation when eCB-CB1 signaling is enhanced by the FAAH inhibitor PF3845 (Fig. 4). Though it is well-reported that eCB activation affects network activities, literature disagrees with the directionality of the regulation, with some reporting a net gain (Pava et al., 2014) in up-state and some others a net loss (Fortin and Levine, 2007; Lafourcade et al., 2007). Since CB1 activation suppresses synaptic vesicle release at both excitatory and inhibitory synapses, these divergent results likely reflect the summation of eCB actions on network activities mediated through inhibitory and excitatory synapses, which may vary based on the specific E/I-balance in the network model. This indicates that the enhanced endocannabinoid tone during BCC-induced network hyperexcitation (Fig. 2) could serve to stabilize network activities and to prevent runaway excitation.

Additionally, we show that CB1 inactivation strongly and transiently suppresses baseline network activities in naïve and downscaled cells, but the suppression is attenuated after downscaling (Fig. 5). This agrees with previous findings that eCB inhibition disrupts network oscillation and sleep stability (Martin et al., 2022; Pava et al., 2014; Pava et al., 2016). The diminished effect of CB1 inactivation after downscaling could be due to BCC masking the disinhibition of GABA release, or due to activity-dependent regulation and complex interaction of CB1-downstream targets suggested in mechanistic models of network oscillation (Compte et al., 2003; Neske, 2015), such as GABA channels (Crosby et al., 2019; Wenner, 2011) or GIRK channels (Guo and Ikeda, 2004; Zou et al., 2019).

Evidence recently emerged from multiple groups that eCB signaling promotes sleep, particularly by promoting the stability of NREM sleep (Bogathy et al., 2019; Kesner and Lovinger, 2020; Martin et al., 2022; Pava et al., 2016), which has been suggested as a result of CB1 modulation of cortical up-states in similar *in vitro* preparations (Pava et al., 2014). Our current findings provide an additional parallel between network adaptation of hyperexcitation in culture, and the synaptic changes that occur during sleep, and suggest that stabilization of synchronous cortical activity provides the basis for the NREM promoting effects of eCB signaling.

## Acknowledgements

This work was supported by a grant to GHD from the National Institutes of Health (RF1AG068063), and to SLG (R01NS112326).

---

**Extended Figure 5-1.**
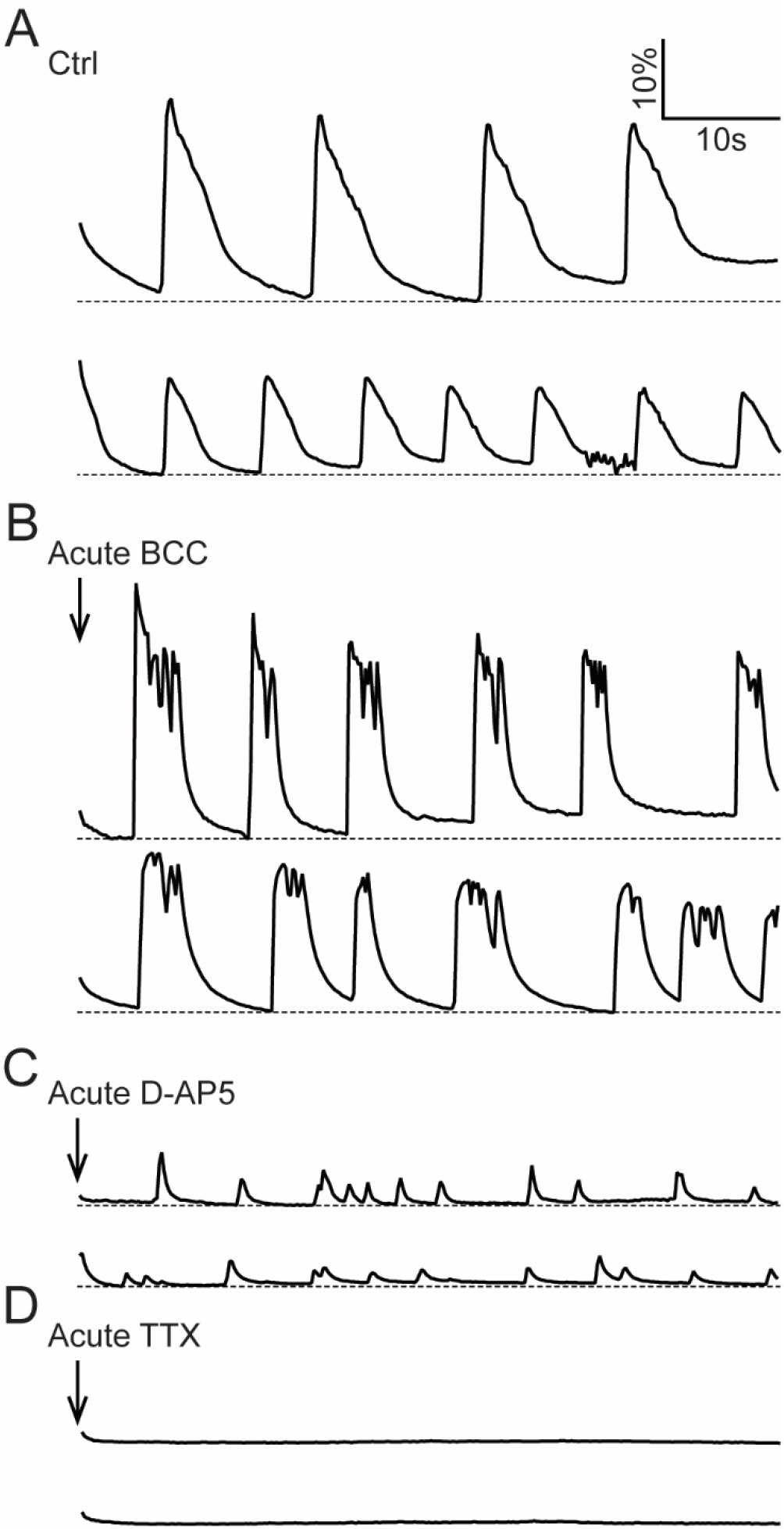
Fluorescence responses of iGluSnFR synchronous events to acute (10min) application of pharmacological treatments: Ctrl **(A)**, 20μM BCC **(B)**, 50μM D-AP5 **(C)** and 1μM TTX **(D)**. Note the similarity to previously reported MEA network activities: glutamatergic events show hyperexcitation to BCC, a GABA antagonist, is dramatically reduced but not eliminated by D-AP5, an NMDA antagonist, and completely abolished by TTX, a voltage-gated sodium channel blocker. Results are presented as change in fluorescence for each recording normalized to its minimum fluorescence (ΔF/F, dotted line represent minimum fluorescence). Scale bar represents 10s (horizontal) and 10% change from minimum fluorescence (vertical). Ctrl data in panel is from the same set of data as in Figure 5.

